# Shared molecular regulation of quiescence in neural and glioma stem cells reveals therapeutic vulnerabilities

**DOI:** 10.1101/2025.01.22.634421

**Authors:** Chandra Choudhury, Matthew Singleton, Stephanie Brauer, Dana Friess, Jessica Hart, Niclas Skarne, Karthik Pullela, Likai Mao, Bryan W Day, Lachlan Harris

## Abstract

Quiescence, a reversible state of cell-cycle arrest, is an adaptive feature of many adult tissue stem cells, including those in the adult brain. In gliomas, brain tumour stem cells that reside in a quiescent state preferentially survive chemotherapy and radiotherapy, highlighting their critical role in therapy resistance and disease progression. To date, it remains unclear whether the molecular programs governing these states are functionally conserved between neural stem cells and brain tumour stem cells. Here, we establish novel *in vitro* models to study quiescence and find that glioma stem cells are markedly more resistant to entering quiescence than neural stem cells, suggesting that glioma stem cell quiescence more closely resembles a slow-cycling phenotype or shallow quiescence. Nonetheless, direct comparison of quiescent neural stem cells and quiescent/slow-cycling glioma stem cells, as they transition towards proliferation, reveals conserved gene expression trajectories, indicating shared molecular mechanisms. Furthermore, we find that pathways influencing quiescence in neural stem cells exert similar effects in glioma stem cells, underscoring the functional parallels between these populations. Finally, we identify that inhibition of TGF-β signalling might provide an avenue to improve current standard-of-care treatments by targeting quiescent glioma stem cells.

**Note on version 2:** This version corrects two figure errors and expands the Methods. The conclusions of are unchanged.

**Images:** In version 1, debris adjacent to the organoid had been digitally removed from the drug-treated image in Figure 6D. The unmodified original is restored here. Images within Figures 3A and 6D have each been processed identically to one another. Scattered cellular debris was present to a similar extent in control and drug-treated samples.

**Data:** The glioblastoma organoid data in Figure 6E have been updated following re-imaging and blinded re-analysis. Reported effect sizes have changed slightly; the conclusions are unchanged. Figure S2 has likewise been updated following re-imaging and re-analysis.

**Methods:** The Methods have been expanded to describe the analysis more fully, including data exclusion criteria, normalisation, and blinding.

Minor changes to figure legends and the main text are not itemised.

## INTRODUCTION

Quiescence is a reversible state of cell-cycle arrest that is an adaptive feature of many adult tissue stem cells, including those found in the central nervous system (*1*). Quiescence is often referred to as a single “G_0_” state; however, cellular quiescence exists on a spectrum between deep and shallow states (*2–4*). Cells residing in shallow quiescence have short activation times as they remain alert and primed for activation. Conversely, cells in deeper states of quiescence require prolonged or stronger growth factor signalling to enter a proliferative state.

In mice, where neural stem cell (NSC) quiescence has been most thoroughly studied among mammals, approximately 95% of adult NSCs are quiescent (*5*). These quiescent NSCs reside in two niches – the hippocampus and the subventricular zone (SVZ) lining the lateral ventricles – where they settle during early postnatal life and then enter quiescence. This initial entry into quiescence (*6*), and the subsequent deepening of quiescence during aging, ensures the long-term maintenance of NSCs throughout life (*7–9*).

Likewise, in brain cancers, emerging evidence suggests that brain cancer stem cells that reside in a quiescent state preferentially survive chemotherapy and radiotherapy. For example, in sonic hedgehog-driven medulloblastoma, quiescent SOX2^+^ cells drive relapse (*10*). Similarly, in *in vitro* models (*11*), genetically engineered mice (*12*) and patient-derived xenografts (*13*) of high-grade glioma, slow-cycling glioma stem cells (GSCs) resist chemoradiation and regrow the tumour once therapy has stopped. Despite the potential therapeutic significance of quiescence during brain cancer progression, it remains unclear whether the molecular programs governing quiescent states are conserved between NSCs and GSCs. While multiple pathways such as BMP signalling (*14*), lipid metabolism (*15*), and immune cell interactions (*16*) are known to maintain NSC quiescence, the therapeutic relevance and tractability of these pathways in brain cancer have received relatively little attention (*17*).

Here, using *in vitro* models of quiescence, coupled with pseudotime trajectory analysis and functional assays, we address these questions. We find that the gene expression trajectories of quiescent NSCs and GSCs as these cells progress towards an active, proliferating state are at least partly conserved between these populations. This conservation includes genes involved in neuronal differentiation, while genes unique to GSC activation include those regulating cellular biosynthesis. Although *in vitro* induction of NSCs and GSCs into a quiescent state demonstrates that GSCs are more resistant to entering deeper states of quiescence, many pathways that either deepen quiescence or promote activation are functionally conserved. Finally, we identify the inhibition of TGF-β signalling as a rational strategy to sensitise quiescent GSCs to standard-of-care therapies and improve glioblastoma treatment.

## RESULTS

### Quiescent/slow-cycling glioma stem cells are abundant in glioblastoma prior to treatment

Despite the potential significance of quiescent GSCs in driving recurrence in glioblastoma, there have been few attempts at quantifying the fraction of GSCs that reside in a quiescent state across a diverse tumour cohort. Serial transplantation studies are the gold standard for identifying GSC populations but because these assays depend on proliferation, they exhibit a selection bias for actively-dividing GSCs. Relying on marker genes to identify GSCs, such as expression of CD133 (*18, 19*), also has limitations as these markers cannot account for intra-tumoural and inter-patient heterogeneity (*20*). Therefore, to address these challenges, we used a genome-wide approach to identify GSCs by quantifying transcriptional diversity with CytoTRACE, where increased diversity indicates a greater stemness potential (*21*).

We applied this tool to publicly available single-nucleus RNA-sequencing (snRNA-seq) data from 10 glioblastoma tumours (*22*) (Figure 1A). To validate the effectiveness of CytoTRACE, we first demonstrated that its scores were, as expected, significantly higher in tumour cells compared to normal brain parenchymal cells (Figure 1B-D). CytoTRACE can produce erroneous results in mixed samples where cell-size differs substantially between cell populations. This phenomenon was evident in our analysis by a population of normal neuronal cells (i.e., large cells) that had unexpectedly high CytoTRACE scores (Figure 1C, D). To improve the fidelity of the stemness ranking, we re-ran CytoTRACE from each tumour sample separately and only on the tumour population to eliminate cell-size effects (Figure 1E, F). Genes associated with high CytoTRACE scores in this analysis were enriched in known stem cell signalling pathways, including *regulation of stem cell differentiation, stem cell development* and *gliogenesis*. These genes also included known GSC markers such as *CD44* (*23*) (Figure 1G). Validating our decision to not rely on single markers but to use a holistic approach, *CD44* and other GSC markers such as *SOX2* (*24*) exhibited a patient-specific enrichment pattern in GSCs (CytoTRACE-high cells) relative to non-stem tumour cells (NSTCs; CytoTRACE-low cells) and a graded enrichment rather than a binary expression pattern (Figure S1). These findings therefore gave us confidence in assigning a GSC identity to individual tumour cells and allowed us to assess the fraction of these cells that were in slow-cycling/quiescent states across the 10 patient tumours.

**Figure 1:**
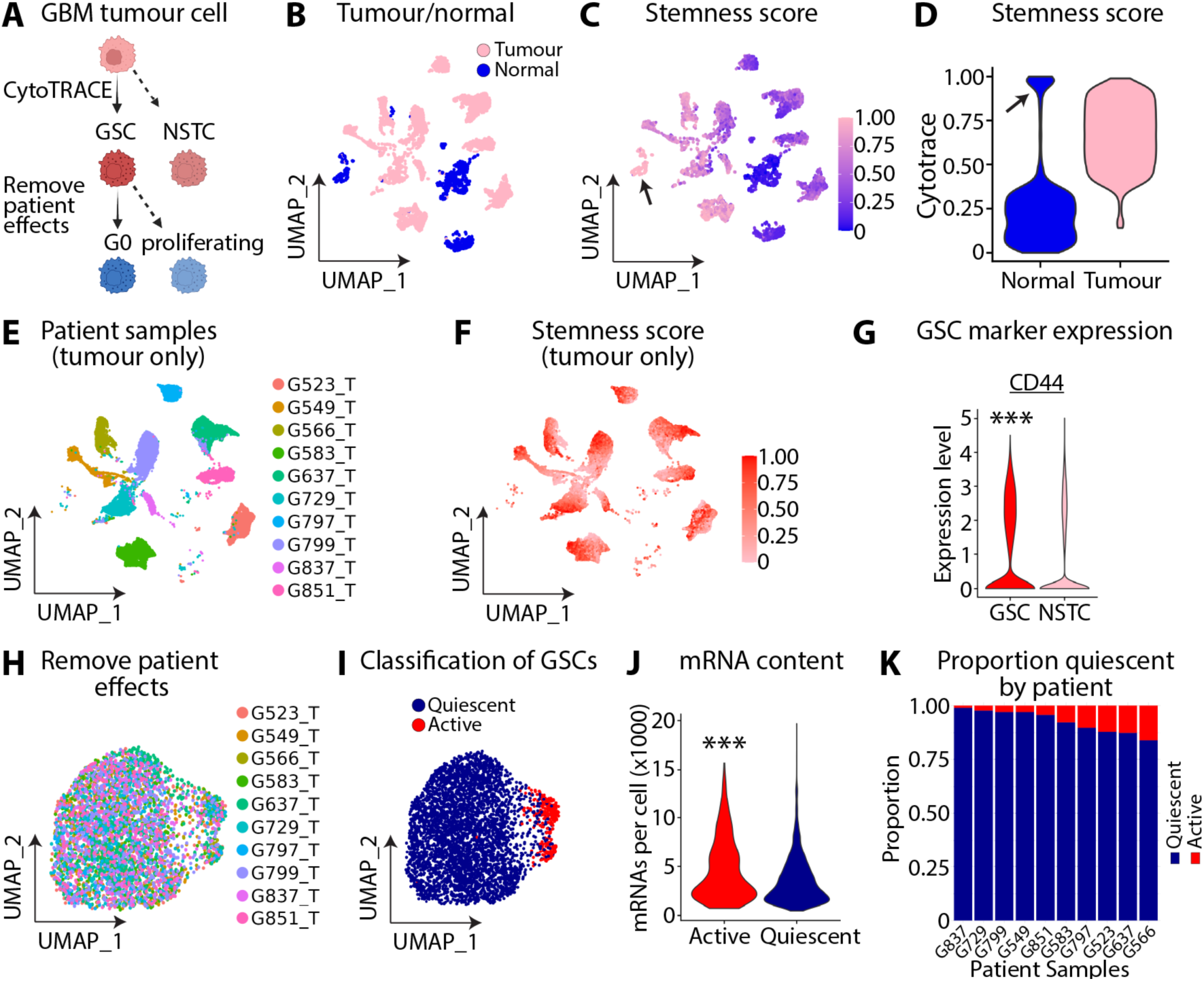
Analysis of snRNA-seq of glioblastoma tumours reveals that quiescent/slow cycling states are predominant in glioblastoma. **(A)** Schematic of classification strategy of quiescent GSCs from glioblastoma tumours. **(B)** UMAP of snRNA-seq data from 10 glioblastoma tumours, coded by tumour-normal status. Dataset from Richards and colleagues (*30*). **(C)** UMAP showing CytoTRACE scores from 10 glioblastoma tumours. Arrow indicates cluster of neuronal cells. **(D)** Violin plot of CytoTRACE scores in tumour and normal cells from 10 glioblastoma tumours. Arrow indicates cluster of neuronal cells. **(E)** UMAP showing malignant cells coded by patient of origin. **(F)** UMAP showing CytoTRACE scores after algorithm run only on malignant cells from each patient independently. **(G)** GSCs (CytoTRACE-high cells) have increased expression of GSC markers than NSTCs (CytoTRACE-low cells). **(H)** UMAP of GSCs only after integration to remove patient effects. **(I)** UMAP showing cluster of active GSCs and quiescent/slow-cycling GSCs. **(J)** Violin plot showing quiescent/slow-cycling GSCs have lower mRNA content than active proliferating GSCs. **(K)** Bar graph showing proportion of GSCs in a quiescent state by tumour sample. Dots in UMAP represent individual cells. Statistics: Mann-Whitney U test in (G, J). \*\*\**P* < 0.001. Abbreviations: GSC – glioma stem cell; NSTC – non-stem tumour cell. Panel A created with BioRender (Agreement number: YQ289ELHTE).

To measure the fraction of GSCs that were quiescent, we reclustered only GSCs and used the anchor-based integration capabilities of the Seurat package (*25*) to identify tumour cells across samples that were related by gene expression profiles (Figure 1H). We identified eight clusters, including two clusters comprised of cells with increased cell-cycle gene expression, which we designated as active cells, while the remaining cells we classified as quiescent (Figure 1I). Quiescent cells have reduced global transcription (*26*). Supporting the validity of the clustering classification, the quiescent clusters had reduced total mRNA levels compared to the proliferating GSCs (Figure 1J). On average, 92% of GSCs resided in a slow-cycling/quiescent state across the 10 tumours (Figure 1K). Independently, we verified these findings by deriving glioblastoma organoids from a surgical resection from a glioblastoma patient. In this heterogenous explant, we found a similarly high percentage of HOPX-(*27*), PTPRZ1-(*28*) and SOX2-expressing cells (*24*) were negative for Ki67 (85-95% of cells) (Figure S2). These data demonstrate that a large fraction of GSCs reside in a slow-cycling or quiescent state prior to chemoradiation and are consistent with previous barcoding studies that suggest treatment primarily selects for pre-existing resistant cell populations within the parental tumour (*13, 29*).

### Quiescent GSCs share activation signatures with healthy neural stem cell populations

The process of stem cells leaving quiescence and proliferating, known as activation, has been well-characterised within the two main neurogenic niches of the adult mouse brain, the hippocampus and SVZ of the lateral ventricles (*17*). We next set out to determine the extent to which the pathways controlling activation in adult NSCs would be conserved in quiescent GSCs. We focussed on the SVZ as this is also thought to be the site of origin for adult high-grade gliomas (*31*). To do this, using single-cell RNA-sequencing (scRNA-seq) data, we ordered NSCs from the mouse SVZ from deep quiescence to an active state using the trajectory inference tool Slingshot (Figure 2A, B) (*32*). We then used tradeSeq (*33*) to identify genes associated with the trajectory (activation signatures) and validated the ordering of these cells by plotting the expression of known markers of deep quiescence, such as *Clusterin* (*34*), and activation, such as *Top2A* (*5*) against pseudotime (Figure 2C). This process was repeated for GSCs derived from the 10 glioblastoma patients (*22*) (Figure 2D-F).

**Figure 2:**
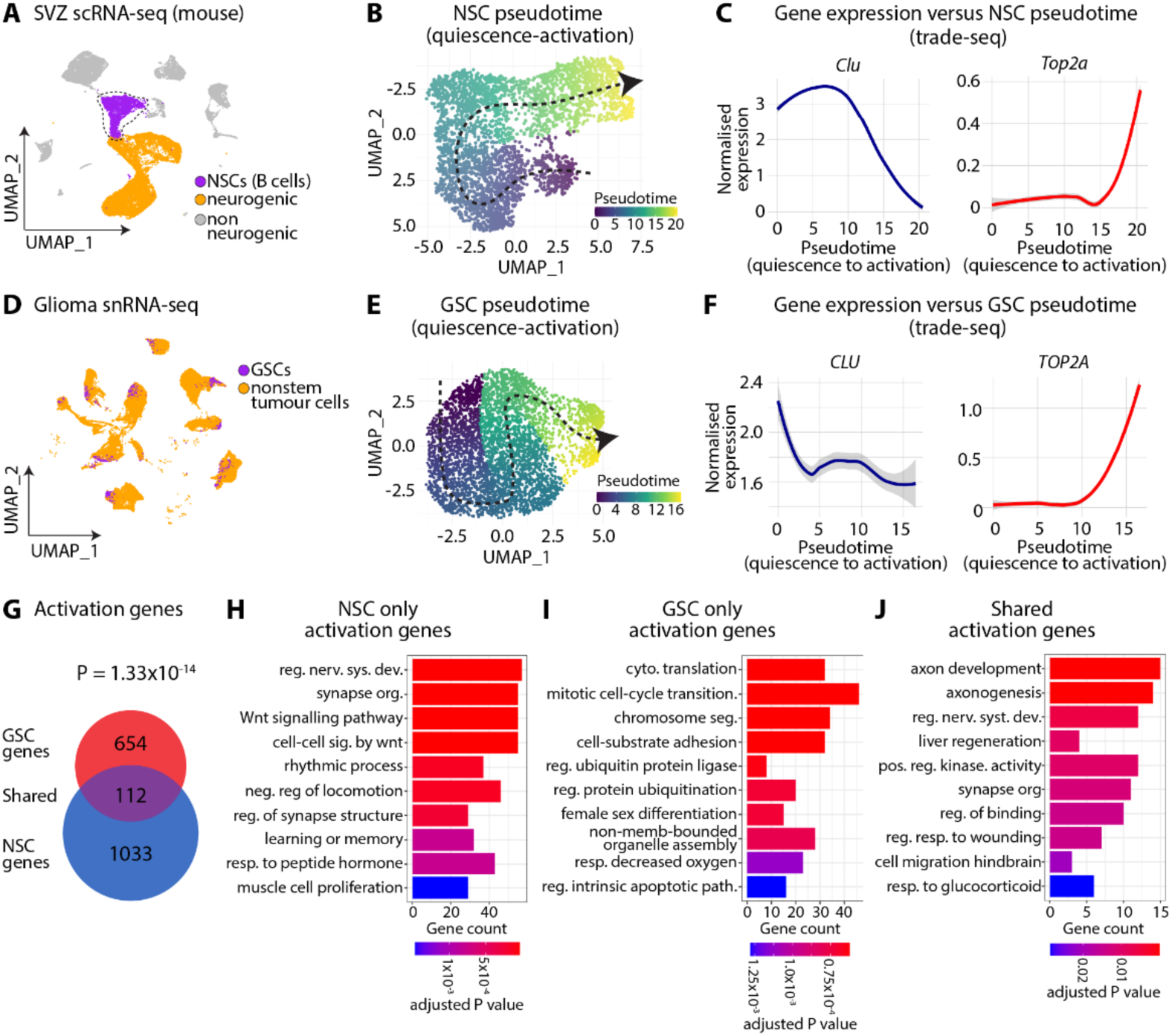
Patient-derived GSCs and adult mouse subventricular NSCs share activation signatures. **(A)** UMAP of scRNA-seq data from mouse subventricular zone showing adult NSCs (B cells), neurogenic cells (type A and C cells), and non-neurogenic cells. Dataset from Cebrian-Silla and colleagues (*31*). **(B)** Pseudotime analysis of NSCs from deep quiescence to an active state using Slingshot. Cells coloured by pseudotime values. **(C)** Expression of quiescence marker *Clusterin* and activation marker *Top2a* against pseudotime. **(D)** UMAP of snRNA-seq data from 10 primary glioblastoma patients showing GSCs, non-stem tumour cells (NSTCs), and non-malignant cells. Dataset from Richards and colleagues (*30*). **(E)** Pseudotime analysis of GSCs from deep quiescence to an active state. Cells coloured by pseudotime values. **(F)** Expression of quiescence marker *CLUSTERIN* and proliferation marker *TOP2A* against pseudotime. **(G)** Overlap of pseudotime-associated genes between NSCs and GSCs. Genes identified with tradeSeq and orthologues with the mouse2human function of homologene. **(H)** Gene ontology analysis of pseudotime genes unique to mouse subventricular NSC activation. **(I)** Gene ontology analysis of pseudotime genes unique to GSC activation. **(J)** Gene ontology analysis of pseudotime genes shared between mouse subventricular NSCs and GSCs. Statistics: Hypergeometric test in (**G**). Arrows in (**B, E**) indicate trajectory from deep quiescence to an active state.

In mouse NSCs, the activation signature comprised 1,403 genes, while in human GSCs 766 genes reached statistical significance (FDR < 0.05; Table S1). Before investigating the overlap between NSCs and GSCs, we used the mouse2human function of homologene to find orthologues for the mouse NSC activation signatures. We found that these signatures were substantially overlapping with the GSCs, approximately 2.1-fold higher than chance alone (Figure 2G). To more closely compare the activation signatures between mouse NSCs and human GSCs, we performed gene ontology of both shared and unique activation genes (Figure 2H-J; Table S2). Shared activation genes were enriched in pathways including *regulation of nervous system development* and *axonogenesis* (Figure 2J), suggesting that both sets of cells are capable of early aspects of neuronal differentiation. Activation genes unique to NSCs were enriched in pathways associated with *Wnt signalling pathway* and *synapse organisation*, consistent with the idea that NSCs are primed to fully differentiate whereas tumour cells are not (Figure 2H) (*35–37*). Finally, activation genes unique to GSCs included those associated with *cytoplasmic translation* (*38*)*, chromosome segregation* (*39*) and *ERK1/ERK2 cascade* (*40*), indicative of a higher rate of biosynthesis and a shallower state of quiescence relative to quiescent NSCs (Figure 2I). Overall, these data demonstrate that while there are distinct activation signatures unique to quiescent mouse NSCs and quiescent human GSCs, many of the core activation processes are conserved.

### Patient-derived GSCs cultured in BMP4 enter a slow-cycling/shallow quiescent state

We next sought to establish *in vitro* models of quiescence in order to test the functional conservation of pathways between NSCs and GSCs. We aimed to develop models that mimicked key features of quiescence – its rapid induction, reversibility and the resistance of quiescent GSCs to current standards of care (*13, 41*). In the first model of quiescence we chose the addition of BMP4, which is the major niche signal that promotes quiescence in mouse NSCs (*14*) and has previously been demonstrated to reduce proliferation in GSC cultures (*11*). To do so, we first derived primary adult mouse SVZ NSCs from C57Bl/6 mice. We then cultured NSCs and GSCs separately in low-levels of growth factors (EGF and FGF) to maintain stemness, before substituting EGF with recombinant human BMP4 (*42*). We titrated BMP4 in the presence of a consistent concentration of FGF and found that administration of BMP4 reliably reduced the proportion of cells expressing Ki67 in a concentration-dependent manner, while retaining expression of the stemness marker SOX2. However, the effect of BMP4 was significantly more modest in GSCs than NSC cultures (Figure 3A-C). The slope of the concentration-response curve was reduced in GSCs approximately 1.6-fold and a comparable reduction in proliferation in the GSC cultures required a BMP4 concentration approximately 160 times higher than that needed for mouse NSCs. These results demonstrate that BMP4 signalling induces a much shallower quiescent state in GSC cultures than in NSC cultures.

**Figure 3:**
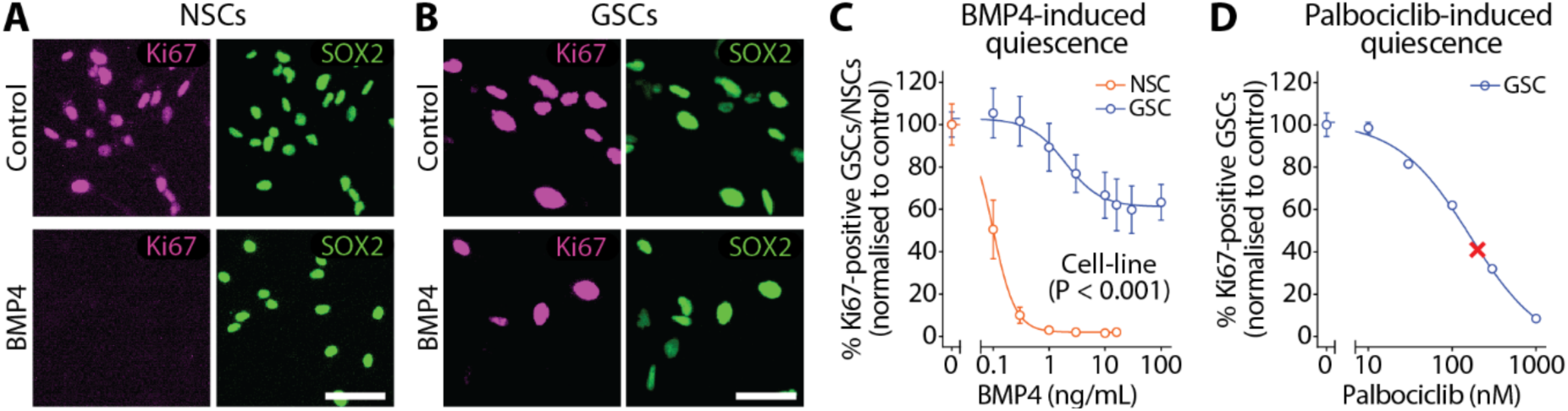
BMP4 and palbociclib treatment induces patient-derived GSCs into shallow versus deep states of quiescence, respectively. **(A)** Treatment with 16 ng/mL BMP4 markedly reduces the proportion of proliferating NSCs (SOX2^+^Ki67^+^) in primary adult mouse SVZ cultures. **(B)** Treatment with 16 ng/mL BMP4 modestly reduces the proportion of proliferating GSCs (SOX2^+^Ki67^+^) in primary patient-derived cultures (QBC395 cell line). **(C)** Concentration-response curve of mouse NSCs and patient-derived GSCs (QBC395 cell line) treated with BMP4, measuring proportion of proliferating stem cells (SOX2^+^Ki67^+^) relative to control media. Data were normalised to mean of control. **(D)** Palbociclib treatment markedly reduces the proportion of proliferating GSCs (SOX2^+^Ki67^+^) relative to control media. Data were normalised to mean of control. GSC graphs in (**C, D**) show mean ± SEM of independent experiments conducted on separate passages (n=3-4). NSC graph in (**C**) shows mean ± SEM of individual cell lines derived from SVZ of different mice (n=4). Scale bar: 15 µm in (**A, B**). Cross in (**D**) represents concentration-response at 200 nM. Statistics in (**C**): two-way ANOVA reporting main effect of cell-line.

### Patient-derived GSCs cultured in low concentrations of CDK4/6 inhibitors enter a deeper quiescent state than in the presence of BMP4

The resistance of the GSC cultures to enter a deeper quiescent state spurred us to also develop a second model. With rare exceptions, such as in retinoblastoma-deficient cancer cells (*43*), entry into G_1_ is governed by the activity of CDK4 and CDK6. We hypothesised that low levels of CDK4/6 inhibitor palbociclib, below concentrations and durations typically required to induce senescence (*44*), would induce a quiescent-like state in GSC cultures, which was more profound than treatment with BMP4. Indeed, we found that the reduction in Ki67 expression was steeply concentration-dependent and that just 200 nM palbociclib caused an approximate 60% reduction in proliferation (Figure 3D).

### BMP4- and palbociclib-mediated quiescence is reversible and confers resistance to radiation

Quiescence is a reversible state of cell-cycle arrest and in glioblastoma confers resistance to standards-of-care. Next, we tested whether the BMP4 and palbociclib models would recapitulate both the reversibility and treatment resistance of the quiescent state. In both experiments, we first induced quiescence using BMP4 or palbociclib for three days (Figure 4A). For the reversal experiment, we performed drug washout and quantified the fraction of Ki67^+^ GSCs at 3, 5 and 7 days afterwards. We found that in both the BMP4 and palbociclib models of quiescence that the proliferation of GSC cultures fully returned to control levels within three days (Figure 4B).

**Figure 4:**
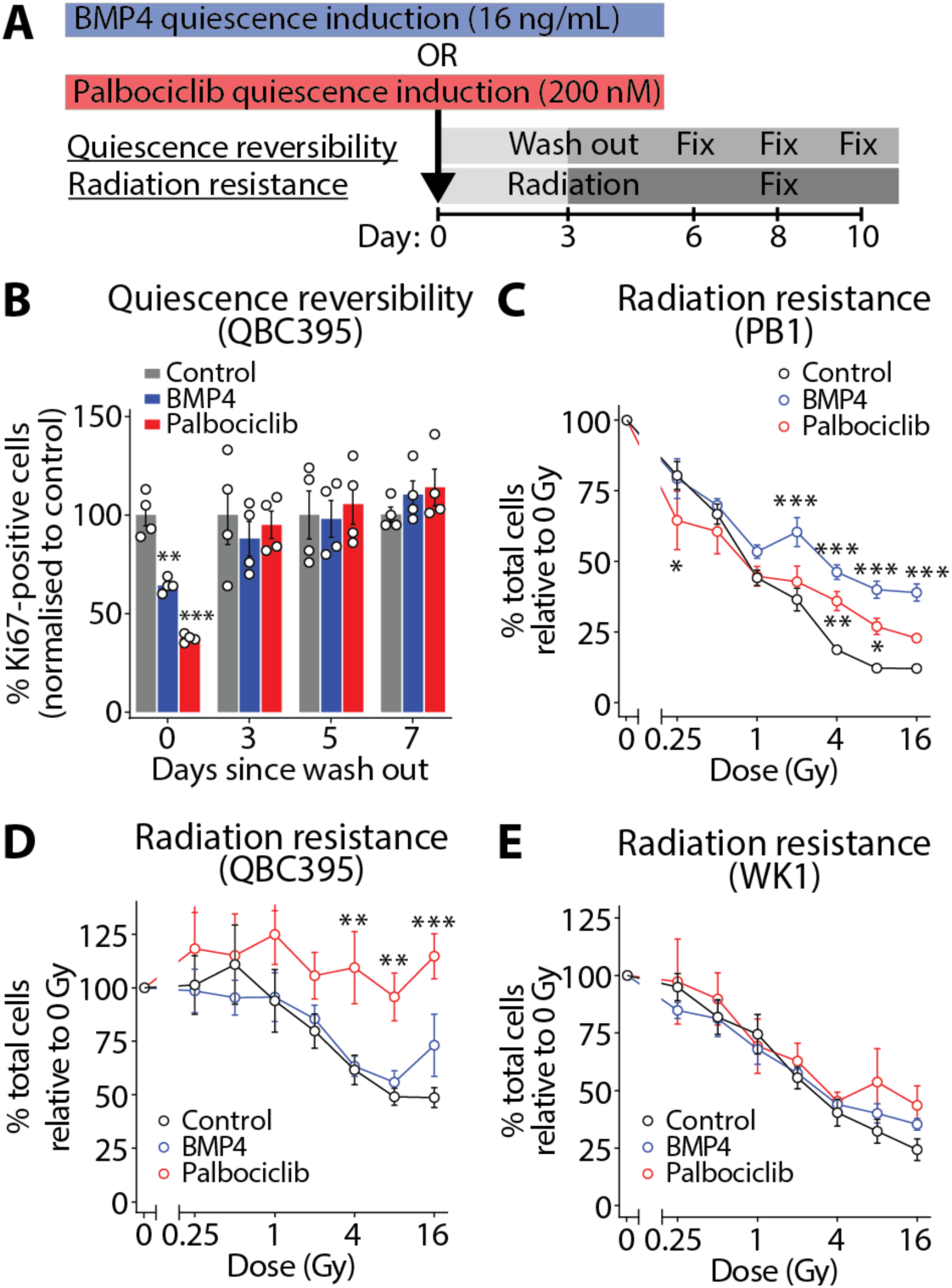
BMP4- and palbociclib-mediated quiescence confers resistance to radiation. **(A)** Schematic outlining different experimental models of quiescence, and the timelines for quiescence reversibility and radiation resistance experiments. **(B)** BMP4- and palbociclib-treated GSCs (QBC395 cell line) returned to control levels of proliferation by three days after wash out (n=3-4). Graph shows percentage of proliferating GSCs (SOX2^+^Ki67^+^) over total GSCs (SOX2^+^). Data were normalised to control media at each time point. **(C)** Effect of radiation treatment on GSC number, relative to 0 Gy control (PB1 cell line, n=4). **(D)** Effect of radiation treatment on GSC number, relative to 0 Gy control (QBC395 cell line, n=9). **(E)** Effect of radiation treatment on GSC number, relative to 0 Gy control (WK1 cell line, n=4). Graphs show mean ± SEM of independent experiments conducted on separate passages. Data in (**C, D, E**) were normalised to 0 Gy control for each independent experiment. Statistics: two-way ANOVA with Holm-Sidak’s multiple comparisons test. \**P* < 0.05, \*\**P* < 0.01, \*\*\**P* < 0.001.

Alternately, after quiescence induction, we treated cultures with radiation and assessed changes in cell-number five days later. We performed this across three primary cell lines with varying levels of radiosensitivity. In the most radiosensitive cell line PB1, both palbociclib- and BMP4-mediated quiescence conferred protection to therapy (Figure 4C), whereas only palbociclib-mediated quiescence provided protection in QBC395 cell line (Figure 4D) and neither conferred protection in the WK1 cell line (Figure 4E). These results demonstrate that in both BMP4 and palbociclib models, quiescent GSCs are relatively protected, though responses are patient- and treatment-specific. Overall, these two models of quiescence recapitulate key features, namely induction, reversibility and resistance, supporting the suitability of these models for screening quiescence-modulating compounds.

### Single-cell RNA-sequencing validates *in vitro* GSC models for screening quiescence-modulating compounds

Up until now, our assessment of quiescence had primarily relied on the absence or presence of Ki67 to indicate quiescent or proliferative states, respectively. However, this approach has limitations—it cannot definitively determine whether cells are simply losing Ki67 expression for unrelated reasons (*8*) or how deep of a quiescent state the cells are in. Thus, before beginning the screen to identify quiescence-modulating compounds in patient-derived GSC cultures, we needed to confirm that our assessment of quiescence, through Ki67 measurements, was reliable. We performed scRNA-seq on low-passage GSCs (4-8 passages) from five patients with glioblastoma in media conditions known to impact quiescence, to varying levels. We grew these cultures in 1) control media, 2) BMP4 to induce quiescence, 3) BMP4 plus palbociclib to induce a deeper state of quiescence and 4) BMP4 and the DYRK1a inhibitor, leucettine L41, which has previously been shown to promote the proliferation of GSCs (*45*).

In total, we sequenced 20,032 cells, with a median of 738 cells per patient, per treatment condition (Figure 5A, B). Using Slingshot, we ordered all the cells from deep quiescence to an active state and validated the ordering of these cells by plotting the expression of known quiescence marker, GFAP (*46*), and activation marker, MKI67 (*47*), against pseudotime (Figure 5C, D). In all GSC cultures, BMP4 treatment caused GSCs to enter earlier stages of pseudotime compared to control cultures, indicative of entering a quiescent state (Figure 5E). Pseudobulk differential gene expression analysis comparing BMP4-treated GSCs with control GSCs revealed an enrichment of pathways including *response to BMP* and *regulation of glial cell differentiation* (Table S3). GSC cultures that were treated with BMP4 plus palbociclib entered even earlier stages of pseudotime than BMP4 alone i.e., moving into deeper states of quiescence (Figure 5E). This also led to additional changes in gene expression associated with *G1/S transition of mitotic cell cycle*, including downregulation of pre-replication licensing genes *MCM2* to *MCM6* (Table S4). Finally, in two of the five patient-derived cell-lines (QBC395 and QBC382), compared to GSCs treated with only BMP4, GSCs that received BMP4 plus leucettine L41 moved into later stages of pseudotime i.e., shallower quiescence (Figure 5E). Together, these findings demonstrate that scRNA-seq reliably captures distinct quiescent states in patient-derived GSC cultures, thus validating our Ki67-based assessment approach and establishing a robust platform for identifying quiescence-modulating compounds.

**Figure 5:**
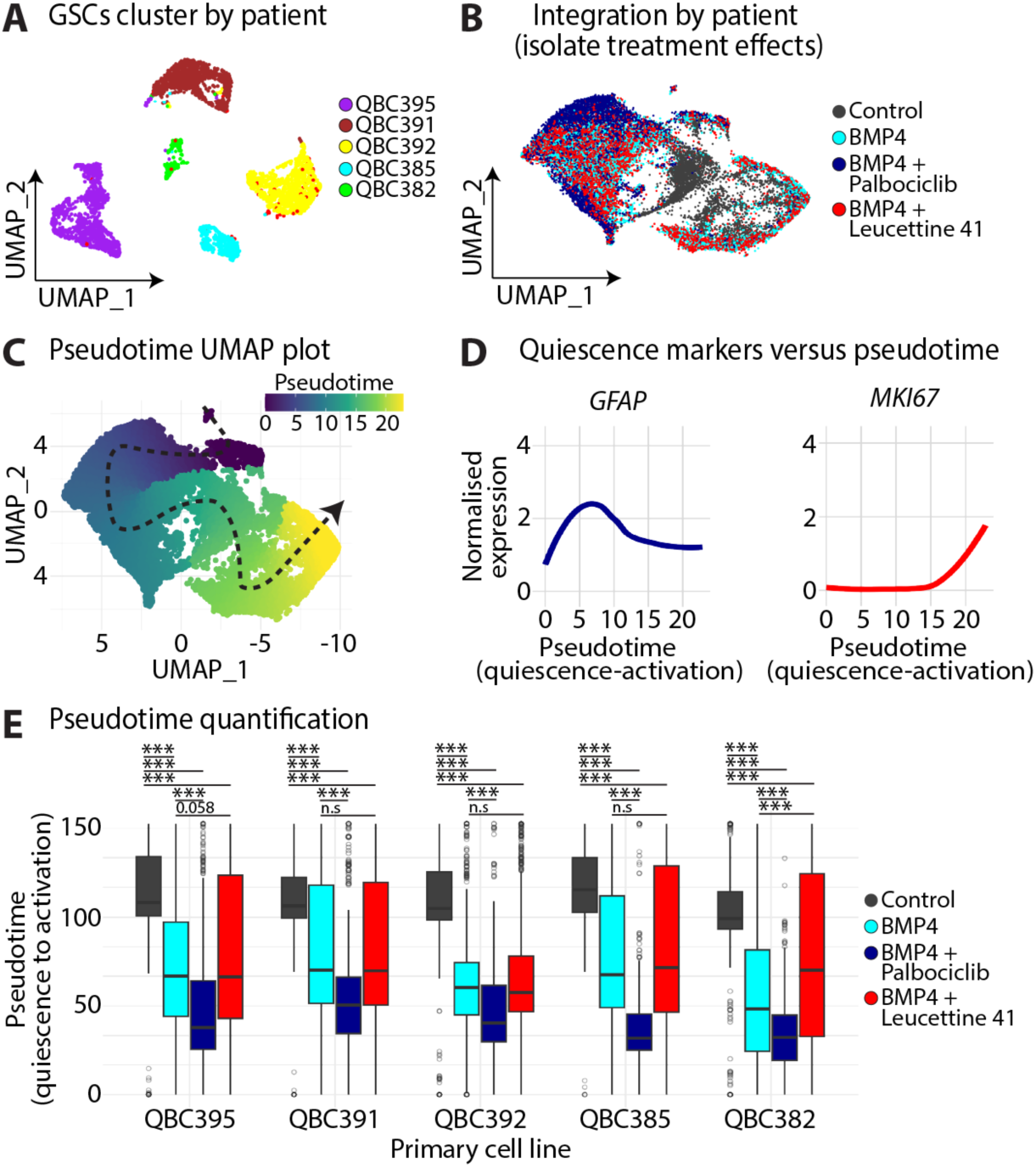
scRNA-seq validates GSC models for screening quiescence-modulating compounds. **(A)** UMAP of scRNA-seq data from five patient-derived GSC lines, each treated with four different culture conditions. Cells cluster by cell-line of origin. **(B)** UMAP of scRNA-seq data integrated by cell-line of origin, revealing biological effects of different treatment conditions. **(C)** Pseudotime analysis of GSCs from deep quiescence to an active state. Cells coloured by pseudotime values, arrow indicates trajectory. **(D)** Expression of quiescence marker *GFAP* and proliferation marker *MKI67* against pseudotime. **(E)** Box plot of change in pseudotime (from deep to shallow quiescence) in response to various treatments. Statistics in (**E**): one-way ANOVA followed by post-hoc Tukey HSD test performed on a per-patient basis. \*\*\**P* < 0.001.

### TGF-β-R1 inhibition drives GSCs out of a quiescent state

Having thoroughly validated the *in vitro* models, we tested a broader range of signalling pathways and targets previously associated with quiescence in mouse NSCs to determine whether they could influence quiescence in GSCs. Specifically, we examined compounds targeting the ubiquitin-ligase HUWE1 (*48, 49*), PTEN (*50*), fatty-acid oxidation (*15*), sonic hedgehog signalling (*51*), interferon signalling (*52*), oleic acid (a ligand for *TLX*) (*53*), and BMP/TGF-β signalling (*14*), which have all been demonstrated to modulate quiescence in adult mouse NSCs in *in vivo* settings. Additionally, we evaluated compounds previously shown to modulate quiescence in patient-derived GSCs including inhibitors against DYRK1A (*45*), PAPD5 (*54*), IDO1 (*55*), CDK4/6 (*56*) and KAT6A (*57*) (Table S5).

We first examined how each compound affected the induction of quiescence in GSC cultures treated with BMP4. We considered a compound to affect quiescence induction if it increased or decreased the proportion of Ki67^+^ GSCs relative to BMP4-only control. As expected, inhibiting BMP4 signalling via the BMP type 1 receptor inhibitor LDN-193189 prevented the induction of quiescence (Figure 6A). Notably, inhibitors targeting TGF-β-R1 (LY-364947), DYRK1a (leucettine L41) (Figure 6A) and histone acetyltransferases (WM-8014; Figure S3) also blocked the induction of quiescence. In contrast, interferon-γ, etomoxir and palbociclib each increased the proportion of cells in a quiescent state, compared to BMP4 alone (Figure S3). To explore the broader applicability of these findings, we then applied inhibitors of TGF-β-R1, BMP, and DYRK1a in our palbociclib-induced model of quiescence. Interestingly, while inhibiting TGF-β-R1 or DYRK1a also prevented the induction of quiescence in this model, inhibiting BMP signalling no longer had an effect (Figure 6B).

**Figure 6:**
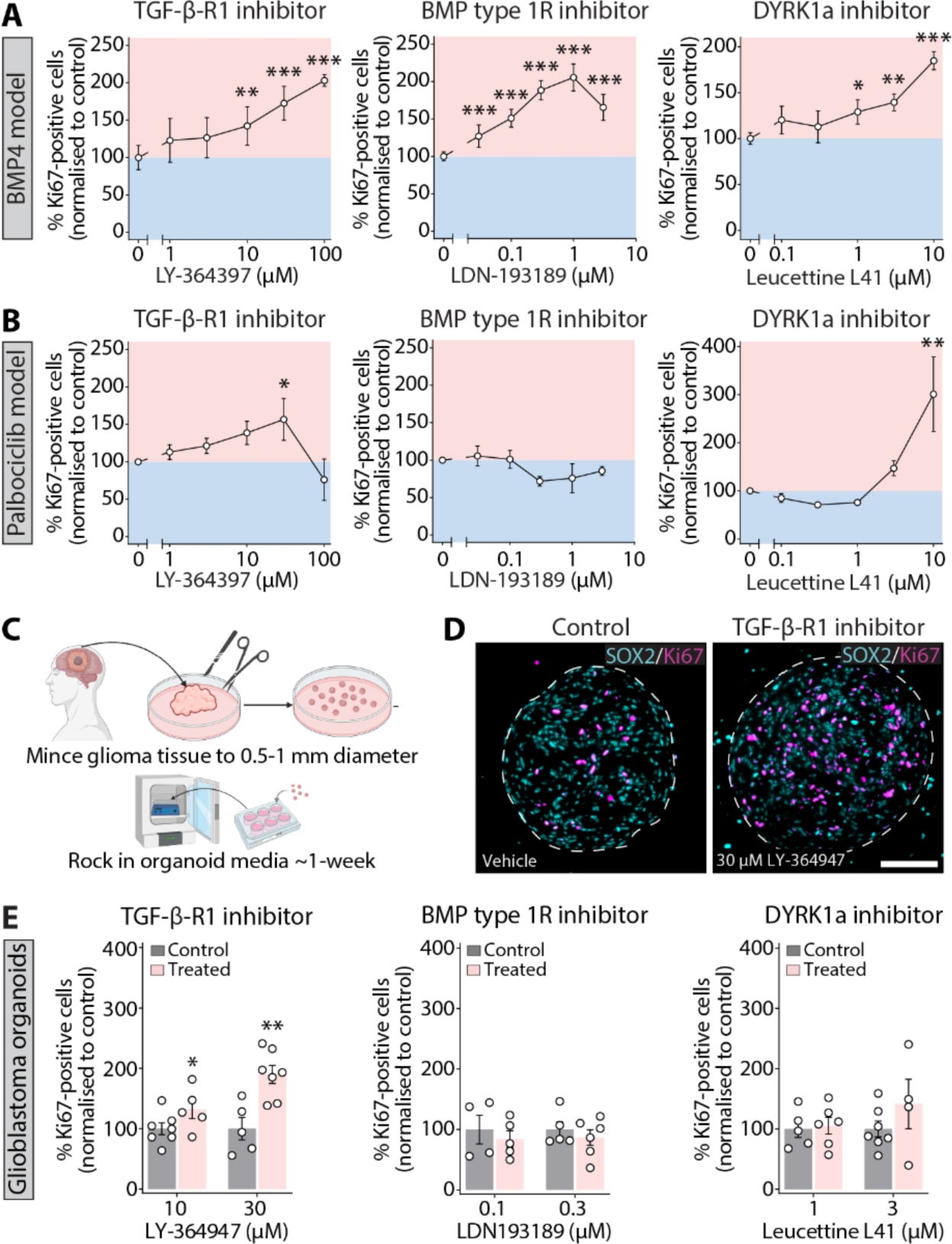
TGF-β-R1 inhibition drives GSCs out of a quiescent state. **(A)** QBC395 cells were cultured in BMP4 quiescence media plus inhibitors for three days. Graphs show percentage of proliferating GSCs (SOX2^+^Ki67^+^) over total GSCs (SOX2^+^). Data were normalised to mean of quiescence media alone. **(B)** QBC395 cells were cultured in palbociclib quiescence media plus inhibitors for three days. Graphs show proportion of proliferating GSCs (SOX2^+^Ki67^+^) over total GSCs (SOX2^+^). Data were first normalised to vehicle control for each independent experiment and then quiescence media alone. **(C)** Schematic of glioblastoma organoid generation. Experiments began within 1-2 weeks of derivation to maximise cellular heterogeneity. Protocol adapted from Jacob and colleagues (*60*). **(D)** Image of glioblastoma organoid treated with 30 µM LY-364397 or DMSO vehicle (control) for three days, stained for SOX2 (cyan) and Ki67 (magenta). Dashed lines demarcate organoid boundary. Scale bar: 100 µm. Scattered cellular debris was observed in both control and drug-treated samples to a similar extent. **(E)** Proportion of proliferating GSCs (SOX2^+^Ki67^+^) over total GSCs (SOX2^+^) after glioblastoma organoids were cultured in the presence of inhibitors. Data were normalised to mean of respective vehicle control. Graphs in (**A, B, E**) show mean ± SEM. Experiments in (**A, B**) represent independent experiments on separate passages (n=3-4). Dots in (**E**) represent individual organoids. Statistics: one-way repeated measures ANOVA with Holm-Sidak’s multiple comparisons test in (**A, B**). Two-way ANOVA in (**E**) with multiple comparisons t-test. \**P* < 0.05, \*\**P* < 0.01, \*\*\**P* < 0.001. Panel C created with BioRender (Agreement number: FY289ELRZ5).

These observations led us to hypothesise that TGF-β ligands might be produced by GSCs themselves to promote a slow-cycling/quiescent state. TGF-β signalling, unlike in many other advanced cancers is intact and overexpressed in glioma (*58*) and has been associated with radioresistance (*59*). To test whether inhibition of TGF-β signalling could reverse quiescence in more complex model systems, patient-derived organoids were generated from glioblastoma surgical resections (Figure 6C) (*60*). Consistent with previous findings, we observed that initiating experiments early maximised organoid heterogeneity, including for example, the retention of endothelial cells (Figure S2), so we began experiments approximately one week after surgery, once the organoids had matured and rounded off (*60*). We observed that the TGF-β-R1 inhibitor LY-364947 at 30 µM caused a 1.85-fold increase in the proliferation of SOX2^+^ GSCs (Figure 6D, E). In contrast, the BMP-signalling inhibitor LDN-193189 (0.1 µM, 0.3 µM) had no effect on the proliferation of SOX2^+^ GSCs (Figure 6E). The DYRK1a inhibitor leucettine L41 (1 µM, 3 µM) also had no effect in the glioblastoma organoids despite positive results in 2D GSC cultures. Together, these results demonstrate that local production of TGF-β ligands – though not BMP ligands – in 2D GSC cultures (Figure 6A, B) and in patient-derived organoids (Figure 6D, E) promotes quiescence and that inhibiting this signalling pathway can promote the activation of quiescent GSCs.

## DISCUSSION

Current therapeutic regimens in glioblastoma, namely the Stupp protocol, primarily target proliferating tumour cells. Consistent with this, there is extensive evidence that tumour recurrence is driven by the preferential survival and reactivation of quiescent GSCs (*13, 41, 61*). Much insight into the regulation of quiescence has been derived from studying adult NSC populations. A recent study suggests these findings can be applied to glioblastoma. For example, increasing FOXG1 alongside WNT signalling successfully induced activation in quiescent NSCs, and this effect was recapitulated in GSCs (*62*). With these findings in mind, we performed a comprehensive comparison of NSCs and GSCs and found substantial transcriptomic similarities during progression from quiescence to an active state, as well as functional similarities in small-scale inhibitor screens. However, we also identified large-scale differences, including that GSCs were significantly more resistant than NSCs to entering quiescence in response to BMP4 treatment, suggesting that in glioma, GSCs may enter a slow-cycling state rather than true quiescence—which is conventionally defined as long-term cell-cycle arrest (*1*). A slow-cycling state may be advantageous to the GSCs, by allowing them to remain poised to re-enter the cell-cycle after treatment cessation (*63*) faster than re-awakening from deeper states of quiescence (*2*).

Most excitingly, due to the number of TGF-β inhibitors in clinical trials for solid cancers (*64, 65*), we found that inhibiting TGF-β-R1 induced the exit from quiescence in GSCs across the multiple cellular models used in our study. These results align with the role of TGF-β signalling in promoting cancer cell dormancy in prostate (*66*) and head and neck squamous cell carcinoma models (*67*). In gliomas, TGF-β is overexpressed compared to normal brain tissue (*68*) and acts as a pleiotropic cytokine regulating both cell proliferation and immune responses (*69*). In the context of GSCs, TGF-β has been implicated in enhancing DNA repair—an effect reversible by pharmacological inhibition (e.g., LY-364947) to increase radiosensitivity (*59*). Our data extend these findings by suggesting that TGF-β inhibition not only sensitises GSCs to radiotherapy due to its role in DNA-repair but may also force these otherwise quiescent or slow-cycling cells into an actively proliferating state, rendering them more vulnerable to standard-of-care treatments. This observation is especially relevant given that the mesenchymal-like state of gliomas, which drives recurrence, is enriched in slow-cycling cells, and is promoted by both immune cell interactions (*70*) and TGF-β signalling (*22*). By disrupting TGF-β-driven quiescence, our findings raise the possibility that combining TGF-β inhibitors with existing therapeutic regimens or novel immunotherapies could shift GSCs out of their protective niche and ultimately improve treatment outcomes.

Alternatively, rather than reactivating quiescent GSCs to increase their sensitivity to chemoradiation in a concurrent setting, another therapeutic approach involves treating with chemoradiation first and then maintaining GSCs in a quiescent state during the adjuvant phase (*71*). Based on our data, following chemoradiation, deepening quiescence by promoting TGF-β signalling or by administration of interferon-γ, etomoxir, or palbociclib could be therapeutically promising. Interferon-γ has notable immunostimulatory effects and combined with its cytostatic properties, this may also be advantageous. The quiescence-promoting activity of interferon-γ has been suggested to depend on Notch activity, a canonical regulator of quiescence in adult NSCs (*72*). Inhibition of fatty-acid oxidation by etomoxir administration has also been shown to significantly reduce the fraction of proliferative adult NSCs *in vivo* (*73*). CDK4/6 inhibitors have primarily been studied in recurrent glioblastoma clinical trials as monotherapies (*74, 75*) or *in vitro*, concurrently with radiation therapy (*76*). However, according to our data, administering these inhibitors shortly after the radiation therapy may be more effective. Modelling approaches to compare the relative benefits of reactivating quiescent GSCs during chemoradiation versus preventing reactivation by maintaining quiescence after chemoradiation will provide guidance to the more clinically-effective approach.

Overall, our findings show that while GSC quiescence more closely resembles a slow-cycling state than canonical quiescence, it shares key molecular programs with quiescent NSCs—and can be targeted therapeutically via TGF-β inhibition, providing a potential avenue to increase the effectiveness of standard glioma therapy.

## AUTHOR CONTRIBUTIONS

Conceptualization, L.H.;

Data Curation, L.H.;

Formal Analysis, C.C., M.S., S.B., J.H., L.M., L.H.;

Investigation, C.C., M.S., S.B., D.F., J.H., N.S., K.P., L.H.;

Visualization, C.C., L.H.;

Methodology, B.W.D, L.H., C.C., N.S.;

Writing – Original Draft, L.H., C.C.;

Writing – Review & Editing, all authors;

Supervision, L.H.;

Funding Acquisition, L.H.;

Project Administration, L.H.

## MATERIALS AND METHODS

### Lead contact

Further information and requests for reagents should be directed to and will be fulfilled by the Lead Contact, Lachlan Harris.

### Materials availability

This study did not generate new reagents.

### Data and code availability

The accession number for the raw RNA sequencing data reported in this paper and the code to reproduce the analyses will be provided upon publication in a peer-reviewed journal.

### Ethics statement

Patient-derived glioblastoma stem cell cultures and organoids were obtained through protocols approved by the Human Ethics Committee of the Royal Brisbane & Women’s Hospital (HREC/17/QRBW/577) and QIMRB Human Research and Ethics Committee, approvals P3420 and P3816.

All experimental protocols involving mice were performed in accordance with the Australian code for the care and use of animals for scientific purposes. This study was approved by the QIMR Berghofer Animal Ethics Committee #A2304-602. Throughout the study, mice were housed in standard cages with a 12 h light/dark cycle and *ad libitum* access to food and water.

## METHOD DETAILS

### Cell Culture

#### Patient-derived GSC culture

GSCs were grown as adherent 2D cultures on flasks coated with Matrigel (Corning, NY, USA) in a 5% CO_2_ atmosphere at 37°C. Cell culture medium was KnockOut DMEM/F-12 supplemented with StemPro Neural Supplement, 1X GlutaMAX, 100 U/mL penicillin-streptomycin, 20 ng/mL EGF, and 20 ng/mL FGF (Thermo Fisher Scientific, MA, USA). Patient-derived GSC cultures (WK1, PB1, QBC382, QBC392, QBC385, QBC395, QBC391) were derived from primary glioblastoma patients (*77–79*) (Table S6). GSCs were induced into quiescence by replacing the cell culture medium containing FGF and EGF with FGF and BMP4 (16 ng/mL). The BMP4 was reconstituted in 5 mM HCl, 0.1% BSA (Sigma-Aldrich, MA, USA), a vehicle confirmed to have no effect on proliferation rates. Alternately, quiescence was induced by adding 200 nM palbociclib (MedChemExpress, NJ, USA) dissolved in DMSO to a final concentration of 0.004% DMSO, which was also confirmed to have no effect on cellular proliferation. At experimental endpoints, GSCs were fixed by incubation with 4% paraformaldehyde (in PBS) for 10 min and washed with cold PBS. To investigate quiescence reversibility, media was refreshed every two days. To investigate radiosensitivity, media was refreshed and then GSCs were irradiated using a Gammacell irradiator (Best Theratronics, ON, Canada).

#### Mouse-derived neural stem cell (NSC) culture

Mice (males/females, 4-6 weeks old) were euthanised by cervical dislocation and the brain was removed and placed in cold PBS for 5 min. Tissue from the SVZ (located in the lateral wall of the lateral ventricles) was dissected. The Neural Tissue Dissociation Kit (P) (Miltenyi Biotec, RBK, Germany) was used to dissociate the tissue and establish single cell suspensions. During the enzymatic digestions, we used a 37°C orbital shaker and following the incubations with enzymatic mix 1 and 2, to aid dissociation, we used manual trituration with fire-polished pipettes (*8*).

NSCs were first cultured in ultra-low attachment 24-well plates (Corning), on an orbital shaker (120 rpm) in a 5% CO_2_ atmosphere at 37°C, to form neurospheres. Cell culture medium was the same as the GSCs. The protocol for neurosphere establishment and culturing was adapted from Deshpande and colleagues (*80*). Briefly, once neurospheres began to form (3-4 days post isolation), they were purified to remove dead cells/debris, and 3-4 days following this, neurospheres were dissociated using Accutase (Thermo Fisher Scientific). From this point, NSCs were grown as adherent 2D cultures, under the same conditions as the GSCs. Quiescence was induced using BMP4 as described for GSCs, except for a lower concentration of BMP4 was used (0.1 ng/mL) in order to provide a comparable reduction in proliferation. At experimental endpoints, NSCs were fixed by incubation with 4% paraformaldehyde (in PBS) for 10 min and washed with cold PBS.

#### Patient-derived glioblastoma organoid (GBO) culture

Tumour tissues were collected from a glioblastoma patient to produce glioblastoma organoids (GBO; patient QBC476, Table S6). GBOs were derived by mincing tissue to 0.5-1 mm diameter pieces, followed by red blood cell lysis, washing in H+GPSA medium, before transferring to GBO medium and incubating at 120 rpm on an orbital shaker (*60*). The GBO medium was 50% DMEM/F12 and 50% Neurobasal supplemented with 1X GlutaMAX, 1X MEM-NEAA, 100 U/mL penicillin-streptomycin, 1X N-2 supplement, 1X B-27 without vitamin A supplement, 55 µM 2-mercaptoethanol (Thermo Fisher Scientific), and 2.7 μg/mL human insulin (Sigma-Aldrich). After shaking for 5-10 days following tumour resection, the GBOs rounded off and ranged in size from 100-600 µm in diameter. Each GBO was placed in a separate well of an ultra-low attachment 24-well plate (Corning) in a final volume of 800 µL GBO medium for drug treatment. Any GBOs that did not round off or significantly darkened, indicating a necrotic core, were discarded. After drug treatment, GBOs were fixed by rocking with 4% paraformaldehyde (in PBS) for 20 min and washed once with cold PBS and twice with 70% ethanol. GBOs were embedded in 2.5% low melting point agarose (Promega, NSW, Australia) and paraffin-embedded and sectioned (to 4 µm). One section per agarose block was subjected to H&E staining to enable visualisation of the sample under a microscope.

### Immunofluorescence staining

To immunostain adherent GSC and NSC cultures, these were blocked in 2% normal donkey serum (in 0.2% Triton X-100 in PBS) for 30 min, with rocking. Cells were then rocked with primary antibodies (anti-Ki67 – 1:200, anti-SOX2 – 1:800) diluted in blocking buffer for 2 h, washed 3 times with PBS for 5 min, and then rocked with fluorescent secondary antibodies for 30 min. Cells were washed again and then rocked with 4′,6-diamidino-2-phenylindole (DAPI, Thermo Fisher Scientific) for 10 min, before one final PBS wash. For experiments where only DAPI-staining was required, cells were permeabilised via a 30 min incubation with blocking buffer, washed once, and then stained with DAPI for 10 min.

To immunostain organoid sections, a PAP pen (Daido Sangyo Co. Ltd., Tokyo, Japan) was used to draw a hydrophobic circle around the slide-mounted GBO. The same immunofluorescence staining method was followed, as above, with the following differences: primary antibody incubation was performed overnight at 4°C, and slides were DAPI counterstained for 30 min. Primary antibody dilutions were: anti-Ki67 – 1:100, anti-SOX2 – 1:400, anti-HOPX – 1:400, anti-PTPRZ1 – 1:400, and anti-CD31 – 1:200. After the final PBS wash, slides were mounted in mounting medium (Dako, CA, USA) and coverslipped.

### Imaging and analysis

To analyse adherent GSC and NSCs stained for SOX2, Ki67 and DAPI, fixed cultures were imaged using an EVOS FL Auto 2 Imaging System (Thermo Fisher Scientific) using the 10X objective. A total of 18 fields of view, or 27-36 for controls, (9 from each replicate well) were captured. CellProfiler (Broad Institute, MA, USA) was used to establish an automated pipeline for cell counting where each field of view was analysed individually (*81*). Cells stained with only fluorescent secondary antibodies and DAPI were used to set the detection thresholds. The proportion of proliferating GSCs (SOX2^+^Ki67^+^ / total SOX2^+^ cells) were quantified. We observed substantial differences in the proportion of proliferating GSCs in areas of decreased or increased cell density. To control for this, we excluded any fields of view that were more than two standard deviations away, in their number of cells or proportion of proliferating GSCs, from the mean of that well. Any fields of view with dust particles or objective lens reflections were also excluded as they impacted the cell count. Finally, the remaining fields of view from the replicate wells were combined and averaged and this value was used in the final analysis for each independent experiment. As the quantification pipeline was automated, we were not blinded to the analysis.

To image and analyse GBOs the entirety of each GBO (approximately 100-600 µM in diameter) was imaged using the 20X objective on the EVOS FL Auto 2 Imaging System. GBOs that were larger than the field of view were imaged via multiple locations and the images were stitched together. To perform cell counts, between 160-200 DAPI^+^ cells were counted, starting in the upper left quadrant, to identify individual cells. Approximately 30% of the GBO was counted. After this, SOX2 expression was visualised. Only SOX2^+^ cells were considered GSCs. Per organoid, a minimum of 150 GSCs were counted, and they were classified as proliferating or quiescent based on the presence or absence of Ki67 expression, respectively. Any smaller GBOs (with a total of 80-250 cells) were counted in their entirety. Any GBOs that did not round off, degraded into debris, or were grossly misshapen were excluded from the analysis. Quantification was performed blinded to drug-treatment.

### Single-cell RNA sequencing

#### Sample Preparation

Patient-derived cultures (QBC382, QBC385, QBC391, QBC392, QBC395) between passages 4-9 were seeded in T75 flasks (Thermo Fisher Scientific) at 4-6 x 10^5^ cells per flask. The subsequent day the cells were washed with PBS and medium was refreshed with control medium containing FGF and EGF, BMP4 quiescence medium (minus EGF + BMP4 16 ng/mL), BMP4 quiescence medium with palbociclib (200 nM) or BMP4 medium plus leucettine L41 (1 µM). After three days, cells were lifted using Accutase (Thermo Fisher Scientific), spun down, supernatant removed, and then washed twice with ice-cold 0.04% BSA (in PBS). The cells were gently resuspended using a wide-bore pipette in 100 µL ice-cold 0.04% BSA (in PBS), counted, and stored on ice. Within each treatment condition, the five cell lines were combined in equal numbers for a final concentration of 1,000 cells/µL. The DNBelab C Series High-throughput Single-cell RNA Library Preparation Set V2.0 (MGI, QLD, Australia) was used to perform the single-cell RNA sequencing, according to the manufacturer’s protocol. Per treatment condition, 10,000 cells were loaded into the scRNA chip for a target recovery of 5,000 sequenced cells.

#### Sequencing, mapping and scSplit

The DNBelab C Series High-Throughput scRNA Analysis Software (dnbc4tools) version 2.1.1 (https://github.com/MGI-tech-bioinformatics/DNBelab_C_Series_HT_scRNA-analysis-software) was utilised for single-cell RNA sequencing analysis. The analysis was performed using the command “dnbc4tools rna run” with default options. The BAM file generated by dnbc4tools was processed to call SNPs using FreeBayes version 1.3, following the recommendations provided on the scSplit homepage (https://github.com/jon-xu/scSplit). Subsequently, scSplit version 1.0.9 was employed for demultiplexing pooled scRNA-seq data. To do so, allele count generation was performed where the command “scSplit count” was executed, producing two allele count matrix files (“ref_filtered.csv” and “alt_filtered.csv”). These files were generated using the BAM file and SNPs obtained from the previous step. The “filter_matrix/barcodes.tsv.gz” file produced by dnbc4tools was provided via the “-b” option and a common human SNP list (hg38) from the 1000 Genomes Project was supplied through the “-c” option, as suggested in the scSplit documentation. Secondly, to perform cell demultiplexing the command “scSplit run” was executed using the allele count matrices to assign cells to distinct samples. The option “-n 5” specified the expected number of mixed samples. Finally, to generate sample genotypes the command “scSplit genotype” was used to produce sample genotypes based on the demultiplexing results. This step generated the output VCF file, “scSplit.vcf”.

#### Seurat analysis: Quality control

The four samples comprising the four different treatment groups were read into Seurat (Ver. 4.3.0.1) for analysis (*83, 84*). We excluded GEMs containing nCount_RNA less than 1000, nFeature_RNA less than 600 or higher than 10% mitochondrial content. In total, 20,508 of 23, 408 cells (87.6%) passed quality control. Cells were assigned to distinct patients based on the output of scSplit. Cells that did not clearly belong to one of the five patient samples were excluded as multiplets. The identity of each of the five patient samples (QBC382, QBC385, QBC391, QBC392, QBC395) was confirmed by performing differential gene expression analysis using FindMarkers and matching these marker genes by performing qPCR.

#### Seurat analysis: Clustering, data visualisation and pseudotime ordering

The merged Seurat object was split according to patient of origin and the data transformed using SCTransform workflow, using vst.flavour = “v2”. The data was integrated using default parameters to control for the genetic background of each patient to isolate the effects of culture conditions. FindClusters function was used with a resolution of 0.3. For pseudotime ordering the GSCs were ordered from deep quiescence to an active state using the pseudotime inference tool Slingshot (*32*) and the UMAP coordinates.

#### Differential gene expression analysis

A pseudo-bulk approach was used to perform differential gene expression analysis between treatment groups. The raw counts matrix was extracted using the Seurat::GetAssayData function, specifying the RNA assay and counts slot. The pseudo-bulk profiles were created by aggregating expression across the five patient and four treatment groups, resulting in 20 profiles. The aggregated counts were formatted for downstream analyses in edgeR (*85*). Normalisation was applied to account for differences in library sizes across pseudobulk samples. The dispersion of the data was estimated using the estimateDisp function, which optimises the fit of the model to the dataset. Differentially-expressed genes were determined using false discovery rate (FDR) < 0.05. Likelihood ratio tests were conducted using the glmLRT function to identify differentially expressed genes for each specified contrast. The results were retrieved using the topTags function, which ranked genes by statistical significance and fold change. Functional enrichment using clusterProfiler (*86*) and the function enrichGO was performed to gain insights into the biological processes altered in response to treatment.

#### Analysis of publicly available single-cell RNA-sequencing data

To analyse mouse NSC data we downloaded dataset GSE165554 (*31*) and used the provided Seurat object and associated metadata to subset B cells. After subsetting the data to only include B cells, we renormalised and reclustered the data using the SCTtransform workflow and then ran Slingshot on UMAP coordinates. For the glioblastoma data, we used snRNA-seq from Extended Figure 9D of Richards and colleagues (*30*), downloaded from the Broad Single Cell Portal, which is comprised of data from 10 patients. The data was first normalised using the NormaliseData workflow and tumour-normal classifications were then applied from the cluster annotations in the metadata. To classify GSCs versus non-stem tumour cells we ran CytoTRACE (*21*) on malignant cells from each of the 10 patient samples separately. We considered cells in the highest decile for CytoTRACE to be GSCs and all other cells to be non-stem tumour cells. After making these classifications, we subsetted the data to only contain GSCs and renormalised the data using the SCTransform workflow, integrating by sample, to leave the main source of variation as biological differences between tumour cells. We ran slingshot on the UMAP coordinates.

In both NSC and GSC datasets tradeSeq (*33*) was applied to fit generalised additive models (GAMs) for each gene along pseudotime (focusing on 5,000 variable genes identified through Seurat::FindVariableFeatures), enabling statistical testing (tradeSeq::associationTest) to identify transcripts that showed significant expression changes. Multiple-testing correction (FDR < 0.05) yielded sets of genes with dynamic expression patterns. Functional enrichment using clusterProfiler::enrichGO was performed to gain insights into the biological processes associated with the pseudotime-dependent gene expression changes that were shared between NSC and GSCs or unique to each individual dataset.

## Quantification and Statistical Analysis

### Statistical analysis of cell counts

The statistical testing approach was implemented using GraphPad Prism (version 9.0) or in R (4.2.0). Two-tailed unpaired Student’s t tests were performed when comparing two groups. A one-way ANOVA was used when comparing three or more groups. For experiments involving two independent variables, a two-way ANOVA was performed. Any significant main effect detected by ANOVA was followed by multiple comparisons analysis. Further details of the statistical tests are included in the figure legends.

Biological replicates for GSC culture experiments comprised of independent experiments performed on separate passages. Data were analysed for each cell line separately and replicates pooled together. Biological replicates for NSC experiments were performed on the same experimental day but comprised of four different primary cell lines.

### Antibodies

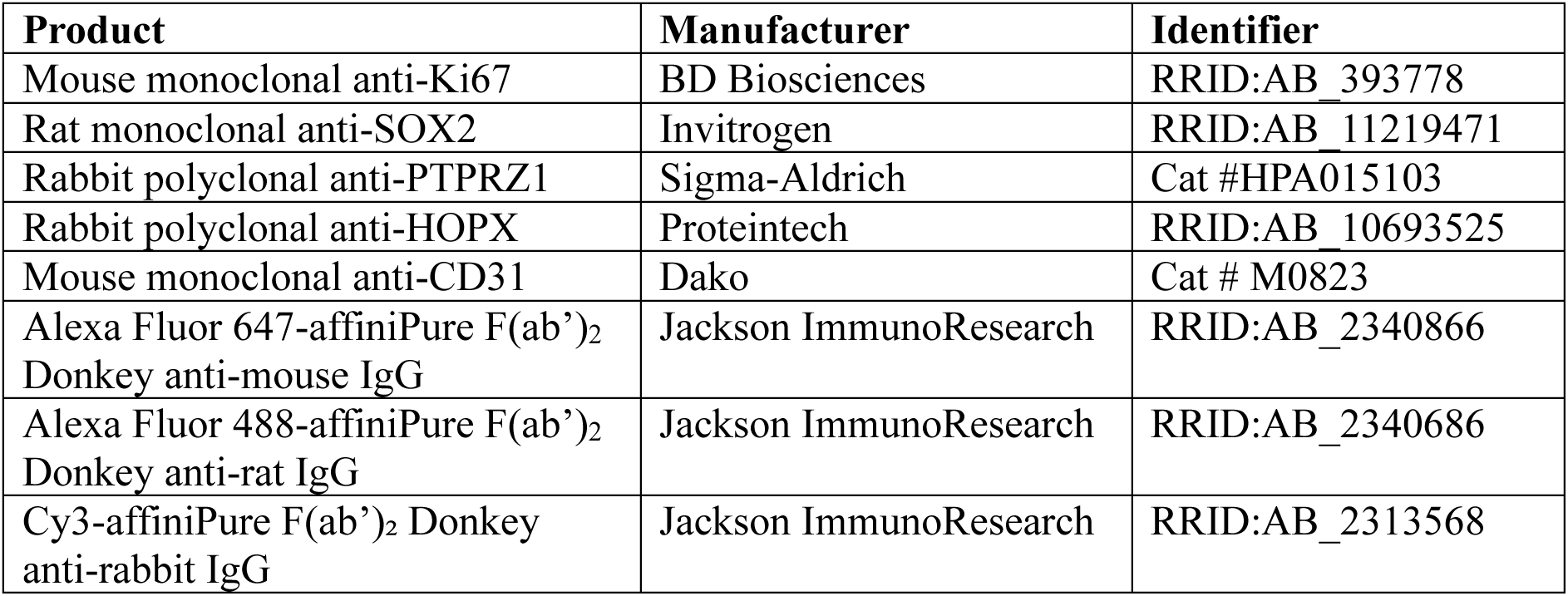

### Critical commercial assays

| Product | Manufacturer | Identifier |
| --- | --- | --- |
| Neural Tissue Dissociation Kit (P) | Miltenyi Biotec | Cat #130-092-628 |
| DNBelab C Series High-throughput<br>Single-cell RNA Library<br>Preparation Set V2.0 | MGI | Cat #940-000519-00 |

### Chemicals, peptides, and recombinant proteins

| Product | Manufacturer | Identifier |
| --- | --- | --- |
| BMP4 | Sigma-Aldrich | Cat #SRP3016 |
| Palbociclib | MedChemExpress | Cat #HY-50767 |
| Temozolomide | Sigma-Aldrich | Cat #T2577 |
| Smoothened agonist | Sigma-Aldrich | Cat #566660 |
| bpV(phen) | Sigma-Aldrich | Cat #203695 |
| Oleic acid | Sigma-Aldrich | Cat # O1257 |
| Leucettine L41 | Cayman Chemical | Cat #29225 |
| BI8622 | MedChemExpress | Cat #HY-120929 |
| BI8626 | MedChemExpress | Cat #HY-120204 |
| LDN193189 (Tetrahydrochloride) | MedChemExpress | Cat #HY-12071A |
| BCH001 | Cayman Chemical | Cat #S8977 |
| SB4 | MedChemExpress | Cat #HY-124697 |
| WM-8014 | Cayman Chemical | Cat #S6655 |
| LY-364947 | MedChemExpress | Cat #HY-13462 |
| Etomoxir | Sigma-Aldrich | Cat #E1905 |
| Epacadostat | MedChemExpress | Cat #HY-15689 |
| Interferon-γ | Gibco | Cat #300-02 |
| DAPI | Thermo Fisher Scientific | Cat #D1306 |

### Key software and algorithms

| Product | Manufacturer | Identifier |
| --- | --- | --- |
| GraphPad Prism 9 | GraphPad Software | RRID:SCR_002798 |
| CellProfiler 4.2.5 | Broad Institute | RRID:SCR_007358 |
| Fiji | Schindelin, *et al.*, 2012 | RRID:SCR_002285 |
| R Project for Statistical Computing 4.2.0 | R Foundation | RRID:SCR_001905 |
| Seurat 4.3.0.1 | Butler, *et al.*, 2018 | RRID:SCR_016341 |
| CytoTRACE 0.3.3 | Gulati, *et al.*, 2020 | RRID:SCR_022828 |
| Slingshot 2.6.0 | Street, *et al.*, 2018 | RRID:SCR_017012 |
| tradeSeq 1.12.0 | Van den Berge, *et al.*, 2020 | RRID:SCR_019238 |
| edgeR 3.40.2 | Robinson, *et al.*, 2010 | RRID:SCR_012802 |
| clusterProfiler 4.6.2 | Wu, *et al.*, 2021 | RRID:SCR_016884 |

## SUPPLEMENTAL INFORMATION

*See Supplemental File and Tables S1-S6*.

## List of Supplemental Tables

**Table S1:** Activation-associated genes in NSCs and GSCs.

**Table S2:** Gene ontology analysis of shared and unique activation genes between NSCs and GSCs.

**Table S3:** Differential expression and gene ontology analysis of BMP4 treated GSCs compared to control GSCs.

**Table S4:** Differential expression and gene ontology analysis of BMP4 plus Palbociclib-treated GSCs compared to control GSCs.

**Table S5:** List of small molecules tested in quiescent GSC models.

**Table S6:** Patient demographics.

## Acknowledgements

We thank Fiona Smith and Courtney Jurd for technical assistance concerning management of brain cancer biobank at QIMR Berghofer and the QIMR Berghofer Histology Facility for their technical assistance in paraffin-embedding and sectioning the GBO samples.

## Supplemental material

**Figure S1:**
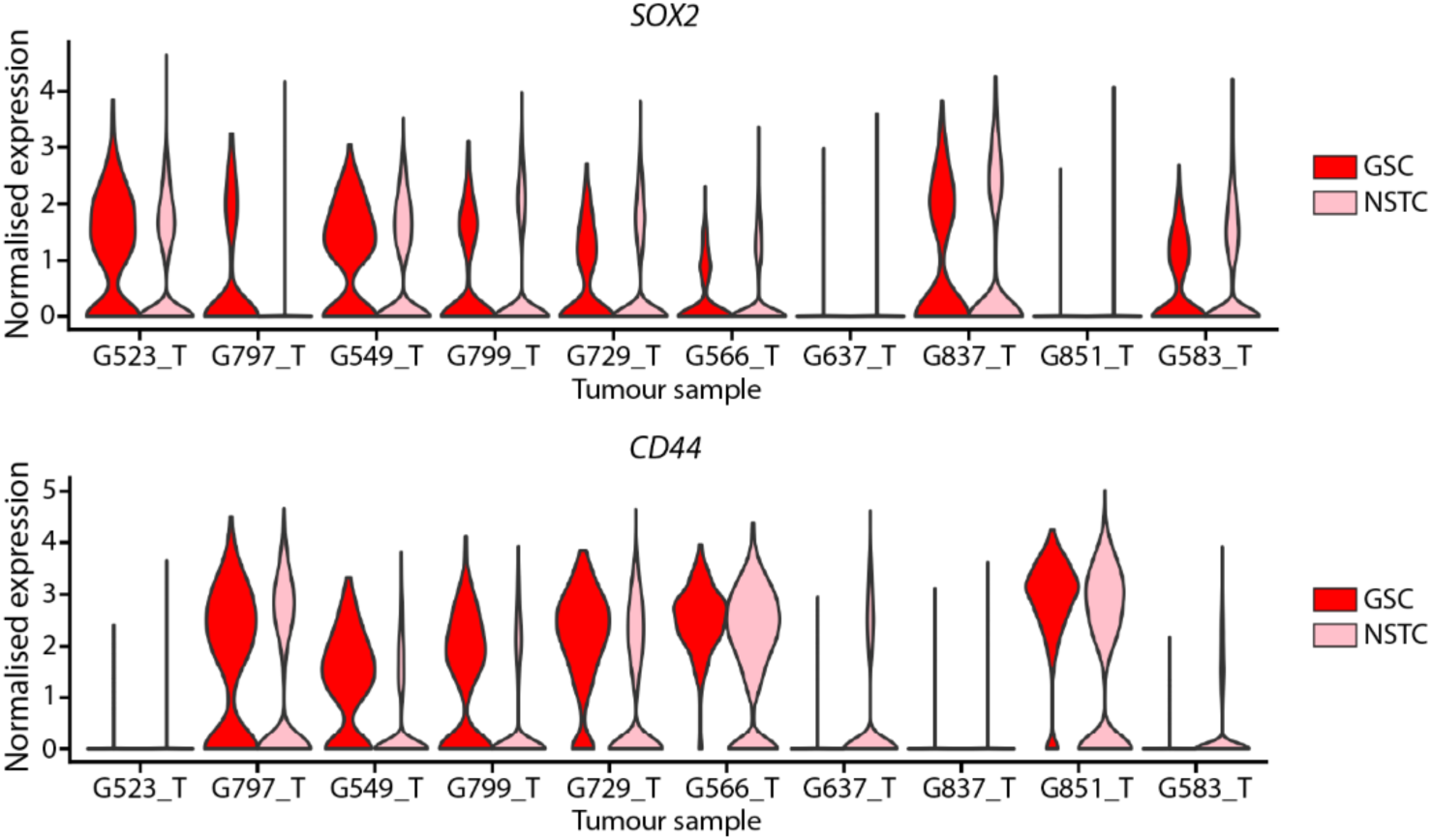
Marker expression in CytoTRACE-high and CytoTRACE-low glioblastoma tumour cells. **(A)** *SOX2* expression in GSCs (CytoTRACE-high) and NSTCs (CytoTRACE-low) glioblastoma tumour cells. **(B)** *CD44* expression in GSCs (CytoTRACE-high) and NSTCs (CytoTRACE-low) glioblastoma tumour cells. Abbreviations: GSC – glioma stem cell; NSTC – non-stem tumour cell.

**Figure S2:**
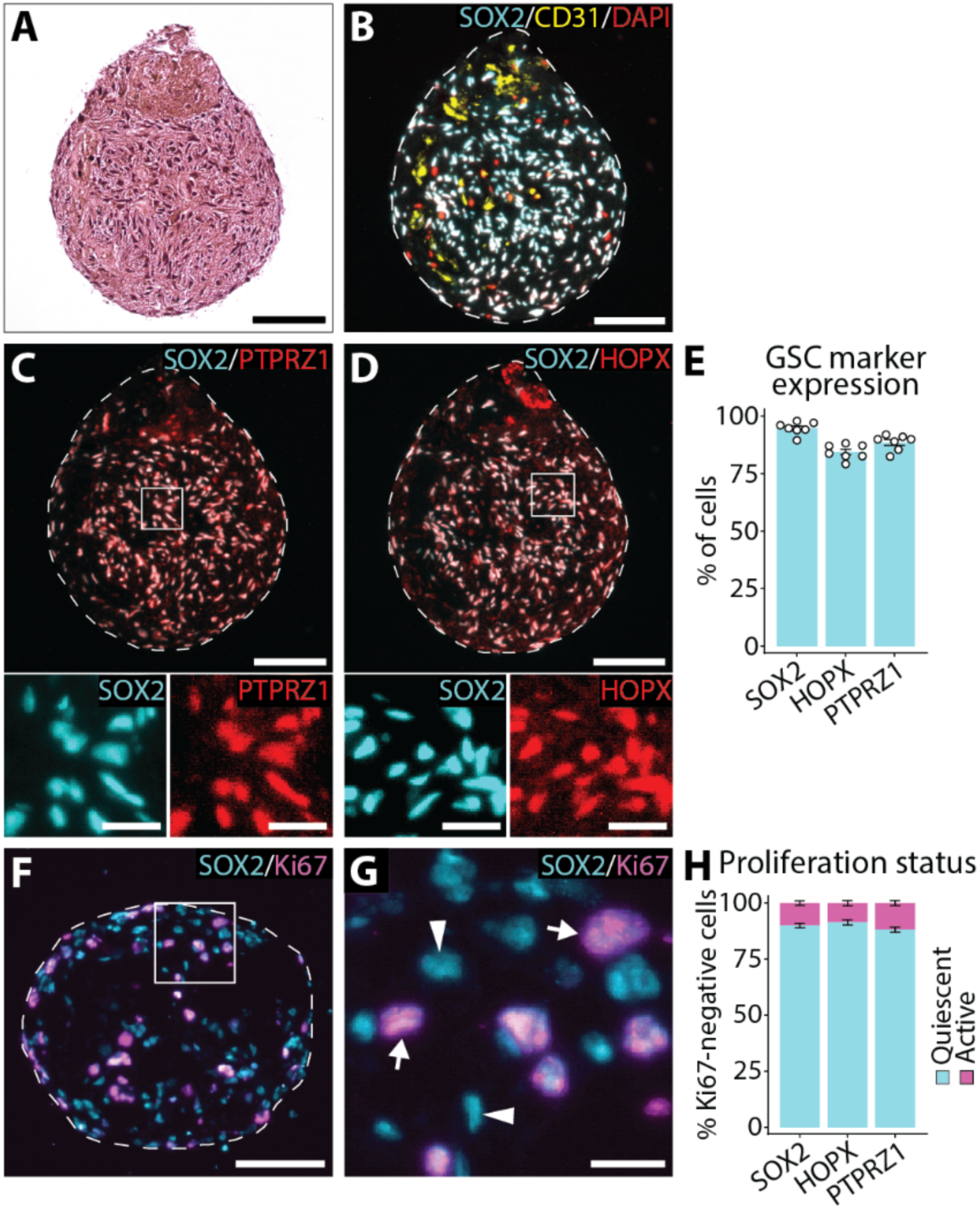
Patient-derived organoids maintain tumour heterogeneity and contain quiescent GSCs. **(A)** H&E stain of GBO. **(B)** Immunofluorescence staining of GBO for SOX2 (cyan), CD31 (yellow) and DAPI (red). **(C)** Immunofluorescence staining of GBO for SOX2 (cyan) and PTPRZ1 (red). **(D)** Immunofluorescence staining of GBO for SOX2 (cyan) and HOPX (red). **(E)** Quantification of the percentage of SOX2, HOPX and PTPRZ1-positive cells. **(F)** Immunofluorescence staining of GBO for SOX2 (cyan) and Ki67 (magenta). **(G)** Inset of (**F**). Arrows indicate Ki67^+^SOX2^+^ cells and arrowheads indicate Ki67^-^SOX2^+^ cells. **(H)** Quantification of the percentage of quiescent (Ki67^-^) and active (Ki67^+^) cells positive for SOX2, HOPX or PTPRZ1. Graphs in (**E, H**) show mean ± SEM, where dots are individual organoids (n=7). Scale bar: 100 μm for whole GBOs in (**C, D, F**) and 20 μm for insets in (**C, D**) and (**G**). Dashed lines demarcate organoid boundary. Abbreviations: GSC – glioma stem cell; GBO – glioblastoma organoid.

**Figure S3:**
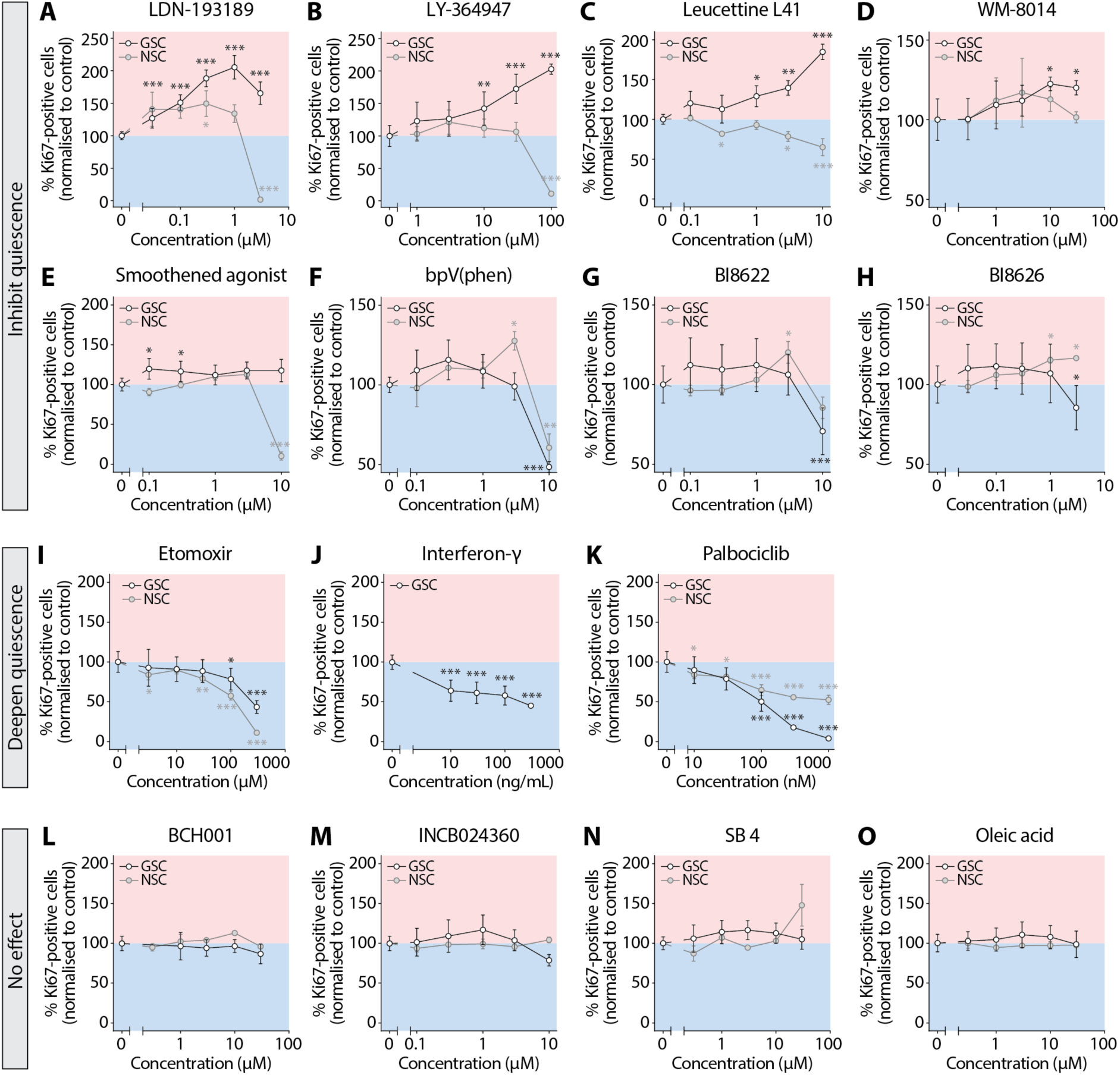
Quiescence modulation in GSC and NSC cultures. (**A-H**) Small molecules that inhibited quiescence in either GSC and/or NSC cultures. (**I-K**) Small molecules that promoted quiescence in both GSC and/or NSC cultures. (**L-O**) Small molecules that had no effect on quiescence in either GSC or NSC cultures. Cells were treated with respective compound and 16 ng/mL BMP4 for three days, and then fixed. Quiescence depth measured as proportion of proliferating stem cells (SOX2^+^Ki67^+^) over total stem cells (SOX2^+^). GSC graphs show mean ± SEM of independent experiments conducted on separate passages (n=3-4) where data were normalised to mean of BMP4 quiescence media alone. NSC graphs show mean ± SEM of individual cell lines derived from SVZ of different mice (n=3-4) where data were normalised to BMP4 quiescence media alone for each cell line. Statistics: one-way repeated measures ANOVA with Holm-Sidak’s multiple comparisons test. \**P* < 0.05, \*\**P* < 0.01, \*\*\**P* < 0.001.

